# Myogenetic oligodeoxynucleotide (myoDN) recovers the differentiation of skeletal muscle myoblasts deteriorated by diabetes mellitus

**DOI:** 10.1101/2021.03.10.434717

**Authors:** Shunichi Nakamura, Shinichi Yonekura, Takeshi Shimosato, Tomohide Takaya

## Abstract

Sarcopenic obesity is a complication of decreased muscle mass and strength associated with obesity, and sarcopenia associated with diabetes mellitus (DM) is a serious risk factor that may result in mortality. Deteriorated differentiation of muscle precursor cells, called myoblasts, in DM patients is considered to be one of the causes of muscle atrophy. We recently developed myogenetic oligodeoxynucleotides (myoDNs), which are 18-base single-strand DNAs that promote myoblast differentiation by targeting nucleolin. Herein, we report the applicability of a myoDN, iSN04, to myoblasts isolated from patients with type 1 and type 2 DM. Myogenesis of DM myoblasts was exacerbated concordantly with a delayed shift of myogenic transcription and induction of interleukins. Analogous phenotypes were reproduced in healthy myoblasts cultured with excessive glucose or palmitic acid, mimicking hyperglycemia or hyperlipidemia. iSN04 treatment recovered the deteriorated differentiation of plural DM myoblasts by downregulating myostatin and interleukin-8. iSN04 also ameliorated the impaired myogenic differentiation induced by glucose or palmitic acid. These results demonstrate that myoDNs can directly facilitate myoblast differentiation in DM patients, making them novel candidates for nucleic acid drugs to treat sarcopenic obesity.

## Introduction

The skeletal muscle is the largest organ associated with metabolism, glucose uptake, thermogenesis, and energy storage. A decrease in muscle mass affects motility, as well as the risk associated with mortality due to chronic diseases such as heart failure (Anker et al., 1997) and cancer (Blauwhoff-Buskermolen et al., 2016). Sarcopenia has been defined as an age-related loss of muscle mass and strength; however, sarcopenia has recently been recognized to arise with obesity, called sarcopenic obesity. Aging and obesity are major risk factors in diabetes mellitus (DM), and the number of patients with DM is increasing worldwide. Muscle wasting is associated with the risk of mortality in patients with DM (Miyake et al., 2019). Therefore, the prevention and treatment of DM-associated sarcopenic obesity is important for public health (Wang et al., 2020). The skeletal muscle is composed of a large number of myofibers, which are multinuclear fused myocytes. Each myofiber has dozens of stem cells, termed satellite cells, between the basal lamina and plasma membrane of the fibers. During myogenesis, satellite cells are activated to myogenic precursor cells, called myoblasts. Following this, myoblasts differentiate into mononuclear myocytes expressing myosin heavy chain (MHC), and mutually fused to form multinuclear myotubes (Dumont et al., 2015). However, the myogenic ability of myoblasts declines with aging or chronic diseases (Fukada, 2018; McCormick and Vasilaki, 2018), which is considered a predisposing factor for amyotrophic disorders.

Both type 1 DM (T1DM) and type 2 DM (T2DM) deteriorate the functions of satellite cells and myoblasts owing to oxidative stress, chronic inflammation, extracellular matrix defects, and transcriptional disorders (D’Souza et al., 2013; Teng and Huang, 2019). In patients with T1DM, the number of satellite cells decreases with the upregulation of the Notch ligand DLL1 (D’Souza et al., 2016). Myoblasts isolated from patients with T2DM show impaired myogenic differentiation with lower miR-23b/27b levels (Henriksen et al., 2017) and autophagy dysregulation (Henriksen et al., 2019). Even after differentiation, myotubes derived from T2DM-patient myoblasts retain an altered myokine secretion distinct from that of non-diabetic myotubes (Ciaraldi et al., 2016). Although the mechanisms underlying the deteriorated function of myoblasts in DM have not been fully elucidated, several factors have been reported to inhibit myogenic differentiation. Co-culture with adipocytes increases interleukin (IL)-6 expression in myoblasts and attenuates their differentiation into myotubes (Seo et al., 2019). High ambient glucose suppresses the myogenesis of myoblasts by increasing the repressive myokine, myostatin, and decreasing myogenic transcription factors, MyoD and myogenin (Grzelkowska-Kowalczyk et al., 2013). Palmatic acid, a saturated fatty acid, blocks myotube formation by downregulating MyoD and myogenin (Saini et al., 2017). These findings demonstrate that diabetic factors including adipokines, glucose, and fatty acids are inhibitory factors for myoblast differentiation.

We recently identified myogenetic oligodeoxynucleotides (myoDNs), which are 18-base single-strand nucleotides that promote myoblast differentiation (Shinji et al., 2021; Nihashi et al., 2021). One of the myoDNs, iSN04, is directly incorporated into myoblasts and serves as an aptamer that physically interacts with nucleolin (Shinji et al., 2021). Nucleolin has been known to target the untranslated region of p53 mRNA to interfere with its translation (Takagi et al., 2005; Chen et al., 2012). In myoblasts, iSN04 antagonizes nucleolin, rescues p53 protein levels, and eventually facilitates myotube formation (Shinji et al., 2021). In this study, we aimed to determine that iSN04 recovers the deteriorated differentiation of myoblasts isolated from patients with DM. This study presents iSN04 as a potential nucleic acid drug targeting myoblasts for the prevention and therapy of sarcopenic obesity.

## Materials and Methods

### Chemicals

The synthetic phosphorothioated iSN04 (5’-AGA TTA GGG TGA GGG TGA-3’) (GeneDesign, Osaka, Japan) was dissolved in endotoxin-free water. Palmitic acid (Wako, Osaka, Japan) was dissolved in chloroform at a concentration of 600 mM (Aguer et al., 2010). An equal volume of endotoxin-free water or chloroform, without the test chemicals, served as negative controls.

### Cell Culture

The human myoblast (hMB) stocks (Lonza, MD, USA) were isolated from healthy subjects (CC-2580) including a 26-year-old (yo) male (H26M; lot 18TL211617), a 35-yo female (H35F; lot 483427), and a 35-yo male (H35M; lot 650386), from patients with T1DM (CC-2900) including an 81-yo male (I81M; lot 211092) and an 89-yo female (I89F; lot 191810), and from patients with T2DM including a 68-yo male (II68M; lot 211384) and an 85-yo female (II85F; lot 219206). The hMBs were maintained in Skeletal Muscle Growth Media-2 (CC-3245; Lonza) as a growth medium for hMBs (hMB-GM). The murine myoblast cell line C2C12 (DS Pharma Biomedical, Osaka, Japan) was maintained in a growth medium for C2C12 cells (C2-GM) consisting of DMEM (Nacalai, Osaka, Japan) with 10% fetal bovine serum and a mixture of 100 units/ml penicillin and 100 μg/ml streptomycin (PS) (Nacalai). hMBs and C2C12 cells were differentiated in a differentiation induction medium (DIM) consisting of DMEM with 2% horse serum (HyClone; GE Healthcare, UT, USA) and PS (Nihashi et al., 2019b; Shinji et al., 2021).

hMB-GM, C2-GM, and DIM with 5.6 mM D-glucose and 19.4 mM mannitol (hMB-GM-NG, C2-GM-NG, and DIM-NG) were used for normal-glucose culture, and those with 25 mM D-glucose (hMB-GM-HG, C2-GM-HG, and DIM-HG) were used for high-glucose culture as previously described (La Sala et al., 2015). In the experiments using high-glucose culture, hMBs were maintained in hMB-GM-HG for a total of six days with passage every three days. The cells were then seeded on fresh dishes and differentiated in DIM-HG for two days. C2C12 cells were maintained in C2-GM-HG for a total of four days with passage every two days. The cells were then seeded on fresh dishes and differentiated in DIM-HG for four days. In the palmitic acid experiments, hMBs were maintained in hMB-GM-NG; then, the cells were seeded on fresh dishes and differentiated in DIM-NG with palmitic acid at an optimal concentration of 200 μM (for H26M) or 600 μM (for H35M) for two days, according to a previous study (Aguer et al., 2010).

All cells were cultured in dishes or plates coated with collagen type I-C (Cellmatrix; Nitta Gelatin, Osaka, Japan) at 37°C with 5% CO2 throughout the experiments.

### Immunocytochemistry

hMBs in hMB-GM (1.5-2.5×10^5^ cells/dish optimized for 70% confluency in each cell stock) or C2C12 cells in C2-GM (10×10^5^ cells/dish) were seeded on 30-mm dishes. The following day, the medium was replaced with DIM containing iSN04 at an optimal concentration of 1 μM (for H26M in hMB-DIM), 3 μM (C2C12 cells), 10 μM (for H26M in GM, H35M, and H85M), or 30 μM (for H35F, I81M, I89F, and II68M). Immunocytochemistry of myoblasts was performed as previously described (Takaya et al., 2017; Nihashi et al., 2019a; Shinji et al., 2021). The myoblasts were fixed with 2% paraformaldehyde, permeabilized with 0.2% Triton X-100, and immunostained with 0.5 μg/ml mouse monoclonal anti-MHC antibody (MF20; R&D Systems, MN, USA) and 1.0 μg/ml rabbit polyclonal anti-nucleolin antibody (ab22758; Abcam, Cambridge, UK). 0.1 μg/ml each of Alexa Fluor 488-conjugated donkey polyclonal anti-mouse IgG antibody and Alexa Fluor 594-conjugated donkey polyclonal anti-rabbit IgG antibody (Jackson ImmunoResearch, PA, USA) were used as secondary antibodies. Cell nuclei were stained with DAPI (Nacalai). Fluorescent images were captured using EVOS FL Auto microscope (AMAFD1000; Thermo Fisher Scientific, MA, USA). The ratio of MHC^+^ cells was defined as the number of nuclei in the MHC^+^ cells divided by the total number of nuclei, and the fusion index was defined as the number of nuclei in the multinuclear MHC^+^ myotubes divided by the total number of nuclei; these were determined using ImageJ software (National Institutes of Health, USA).

### Quantitative Real-time RT-PCR (qPCR)

Total RNA of the myoblasts was isolated using NucleoSpin RNA Plus (Macherey-Nagel, Düren, Germany) and reverse transcribed using ReverTra Ace qPCR RT Master Mix (TOYOBO, Osaka, Japan). qPCR was performed using GoTaq qPCR Master Mix (Promega, WI, USA) with StepOne Real-Time PCR System (Thermo Fisher Scientific). The amount of each transcript was normalized to that of human glyceraldehyde 3-phosphate dehydrogenase gene (*GAPDH*) and murine 18S ribosomal RNA (*Rn18s*). Results are presented as fold-change. The primer sequences are described in Supplementary Tables 1 and 2.

### Statistical Analyses

Results are presented as the mean ± standard error. Statistical comparisons were performed using unpaired two-tailed Student’s *t*-test or multiple comparison test with Tukey-Kramer test following one-way analysis of variance. Statistical significance was set at *p* < 0.05.

## Results

### DM Deteriorates Myoblast Differentiation

The hMBs isolated from healthy subjects (H26M, H35F, and H35M), patients with T1DM (I81M and I89F), and patients with T2DM (II68M and II85F) were cultured in the hMB-GM-NG (Supplementary Figure S1). These hMBs varied in cell size and morphology, but DM-dependent hallmarks were not observed. The hMBs induced myogenic differentiation in DIM-NG, followed by immunostaining for MHC, a terminal differentiation marker of muscle cells. The ratio of MHC^+^ cells and multinuclear myotubes was quantified on days 0, 2, and 4 of differentiation (Supplementary Figure S2). On day 2 (Figure 1), the ratio of MHC^+^ cells of H35M was lower than that of H26M and H35F, indicating the individuality of myogenesis among healthy subjects. I81M differentiated to the same extent as H26M and H35F, but I89F, II68M, and II85F exhibited deteriorated differentiation. In particular, I89F and II85F were exacerbated in myotube formation compared to all healthy subjects. These results indicate that myoblast differentiation is aggravated in patients with DM.

**Figure 1.**
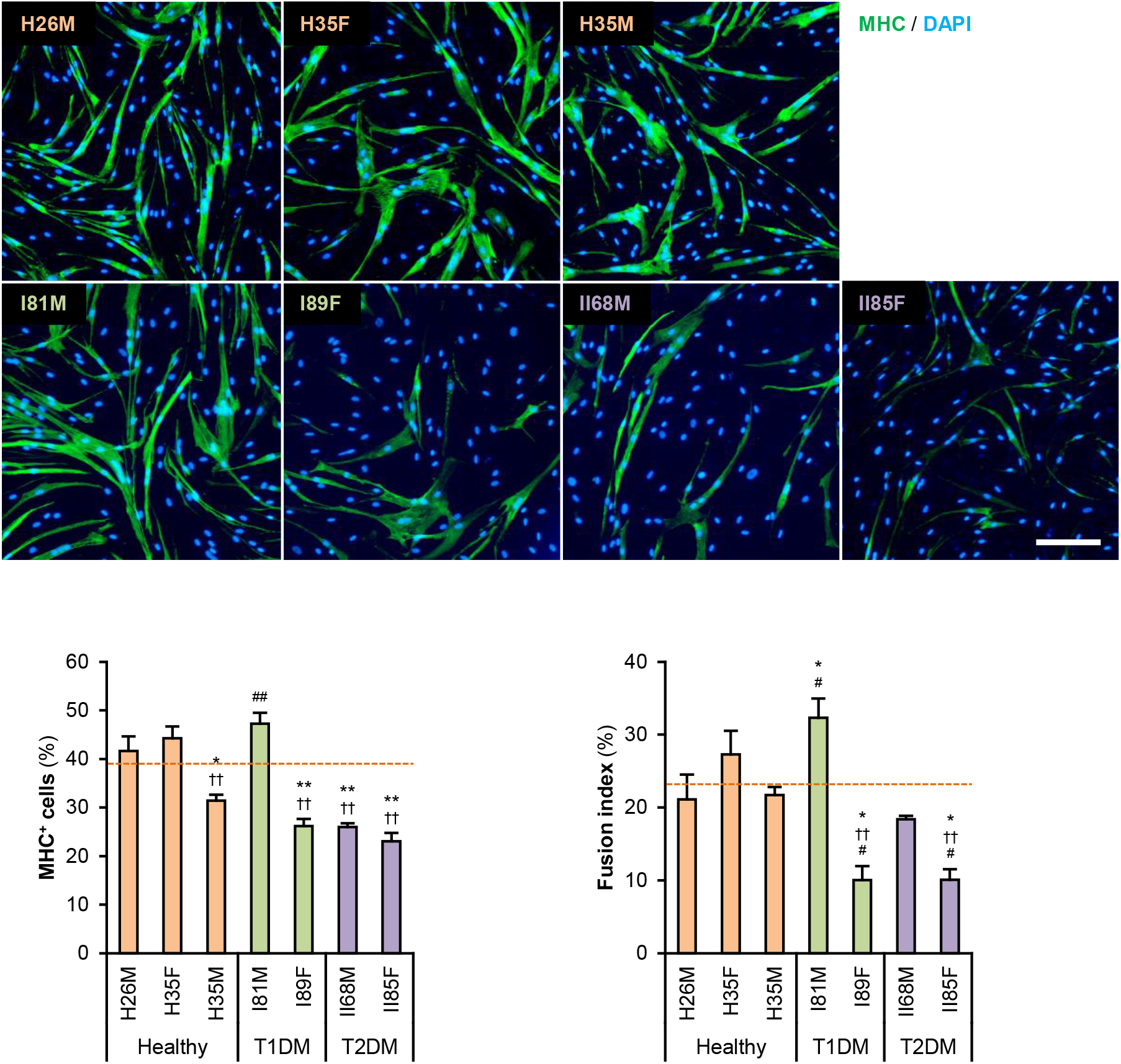
Attenuated myogenic differentiation of DM myoblasts. Representative immunofluorescent images of the hMBs differentiated in DIM-NG for two days. Scale bar, 200 μm. Ratio of MHC^+^ cells and multinuclear myotubes were quantified. Orange dashed lines indicate the mean values of H26M, H35F, and H35M. * *p* < 0.05, ** *p* < 0.01 vs. H26M; †† *p* < 0.01 vs. H35F; # *p* < 0.05, ## *p* < 0.01 vs. H35M (Tukey-Kramer test). *n* = 6.

Gene expression patterns in hMBs were examined using qPCR (Figure 2A). Among undifferentiated myoblast markers, *PAX7* was expressed 2-3 times higher in T2DM myoblasts throughout differentiation, but *PAX3* and *MYF5* were not. A myogenic transcription factor, *MYOD1*, was highly induced in T1DM myoblasts, but a terminal transcription factor, myogenin (*MYOG*), was not. The mRNA levels of embryonic MHC (*MYH3* were not significantly different among hMBs. The transcription levels of these genes frequently vary among patients, which reflects individual differences. During myogenic differentiation, the ratios of Pax7, MyoD, and myogenin are critically important. Proliferating myoblasts express both Pax7 and MyoD, but not myogenin. At the initial stage of differentiation, Pax7 disappears, and MyoD drives myogenin transcription. In terminally differentiated myocytes, MyoD decreases, and myogenin becomes a dominant transcription factor (Dumont et al., 2015). qPCR data indicated that *MYOD1/PAX7* and *MYOG/MYOD1* ratios were lower in T2DM and T1DM myoblasts than those in healthy myoblasts (Figure 2B), demonstrating a delayed shift of myogenic transcription factors in DM patients. This may be one of the reasons for the deteriorated differentiation of DM myoblasts.

**Figure 2.**
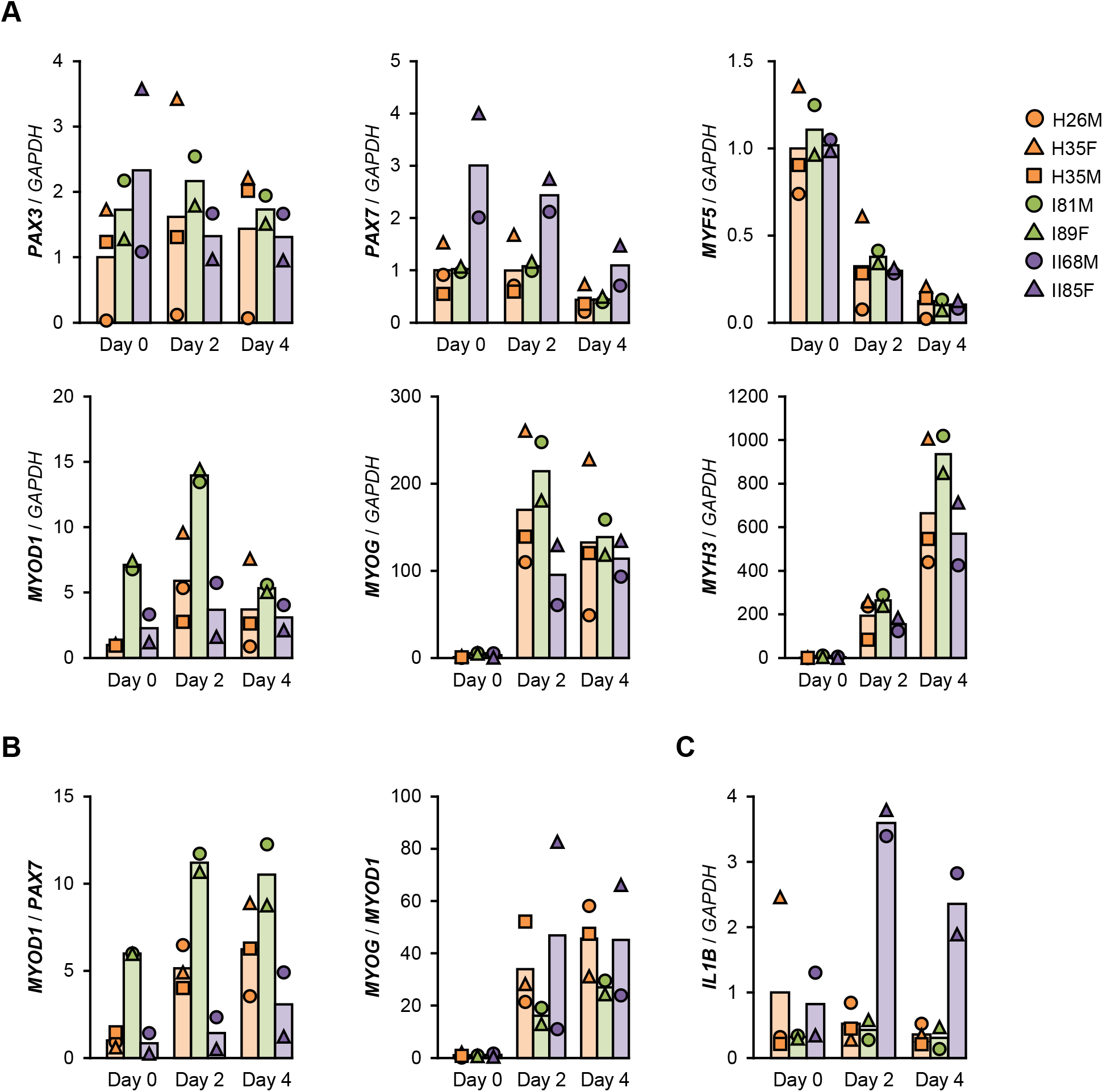
Gene expression patterns altered in DM myoblasts. **(A-C)** qPCR results of gene expression in the hMBs differentiated in DIM-NG on days 0, 2, and 4. Bars indicate mean values of each group. The mean value of healthy myoblasts at day 0 was set to 1.0 for each gene.

### ILs Are Induced in T2DM Myoblasts

The mRNA levels of atrogin-1 (*FBXO32*), MuRF-1 (*TRIM63*), myostatin (*MSTN*), and myostatin receptor (*ACVR2B*), which are involved in ubiquitin-proteasome-mediated muscle atrophy (Bodine et al., 2001; Lokireddy et al., 2011), were not different among the hMBs. In contrast, transcription of the myostatin antagonist, follistatin (*FST*), was flat in T1DM myoblasts during differentiation (Supplementary Figure S3).

Sterol regulatory element-binding proteins (*SREBF1* and *SREBF2*), fatty acid synthase (*FASN*), receptor substrates (*IRS1* and *IRS2*), glucose transporter 4 (*SLC2A4*), mitochondrial carnitine palmitoyltransferase 2 (*CPT2*), and thioredoxin interacting protein (*TXNIP*) are insulin resistance-related factors and involved in differentiation and fatty acid metabolism of muscle cells (Parikh et al., 2007; Kato et al., 2008; Lecomte et al., 2010; Boufroura et al., 2018). However, their mRNA levels were not significantly altered in T2DM myoblasts (Supplementary Figure S4).

T2DM myoblasts have been reported to display abnormal inflammatory responses (Green et al., 2011). Indeed, mRNA levels of *IL1B* were 6-7 times higher in T2DM myoblasts than those in healthy myoblasts on days 2 and 4 (Figure 2C). In contrast, inflammatory factors, NF-κB p50 (*NFKB1* and p65 (*RELA*) subunits, TNF-α (*TNF*), interferon γ (*IFNG*), and *IL6* were not upregulated in T2DM myoblasts. Although *IL8* (*CXCL8*) levels were high in H26M on day 0, T2DM myoblasts exhibited higher *IL8* mRNA levels than those did healthy myoblasts (Supplementary Figure S5). It has been reported that IL-1β inhibits insulin-like growth factor (IGF)-dependent myoblast differentiation (Broussard et al., 2004), and IL-8 is secreted from insulin-resistant myotubes (Bouzakri et al., 2011). Thus, the upregulation of IL-1β and IL-8 potentially impaired the shift in myogenic transcription factors and subsequent differentiation of T2DM myoblasts.

### myoDN Recovers Differentiation of DM Myoblasts

We recently identified the single-strand myogenetic oligodeoxynucleotides (myoDNs) that promote myoblast differentiation by antagonizing nucleolin (Shinji et al., 2021). To assess the applicability of myoDN to DM myoblasts, the hMBs used in this study were treated with iSN04, which exhibits the highest myogenetic activity among the myoDNs. iSN04 significantly facilitated the differentiation and myotube formation of H35F, H35M, I81M, I89F, and II85F (Figure 3). In particular, iSN04 recovered the attenuated differentiation of II85F to almost the same extent as that of healthy myoblasts. iSN04 did not affect the differentiation of H26M in DIM, but significantly promoted myotube formation in hMB-GM (Supplementary Figure S6). In contrast, differentiation of II68M was not altered by iSN04, suggesting the distinct sensitivity or efficacy of iSN04 among individuals. These results indicate that iSN04 is able to recover the deteriorated differentiation of DM myoblasts. qPCR revealed that iSN04 treatment significantly reduced *PAX7* and *MSTN* mRNA levels in II85F, resulting in the recovery transcription of *MYH3* (Figure 4A). An iSN04-dependent decrease in *MSTN* expression was also detected in H35F. Furthermore, iSN04 significantly suppressed the *IL1B* levels in H35F and the *IL8* levels induced in II85F (Figure 4B). These results indicate that iSN04 facilitates the differentiation in both healthy and diabetic myoblasts, in part, by modulating the expression of cytokines including myostatin and ILs.

**Figure 3.**
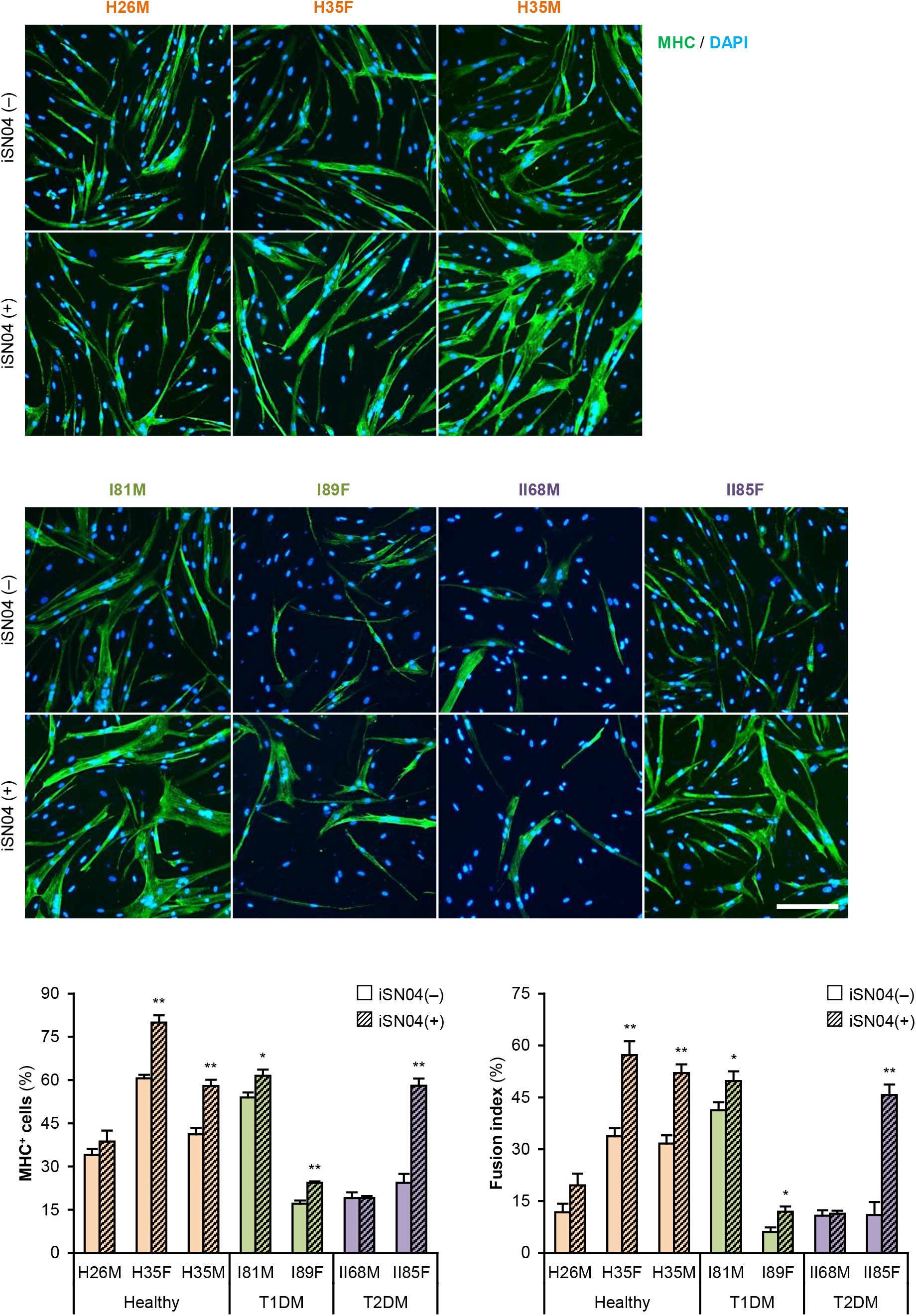
iSN04 recovers the differentiation of DM myoblasts. Representative immunofluorescent images of the hMBs differentiated in DIM-NG with iSN04 for two days. Scale bar, 200 μm. Ratio of MHC^+^ cells and multinuclear myotubes were quantified. * *p* < 0.05, ** *p* < 0.01 vs. iSN04(-) in each group (Student’s *t*-test). *n* = 4-6.

**Figure 4.**
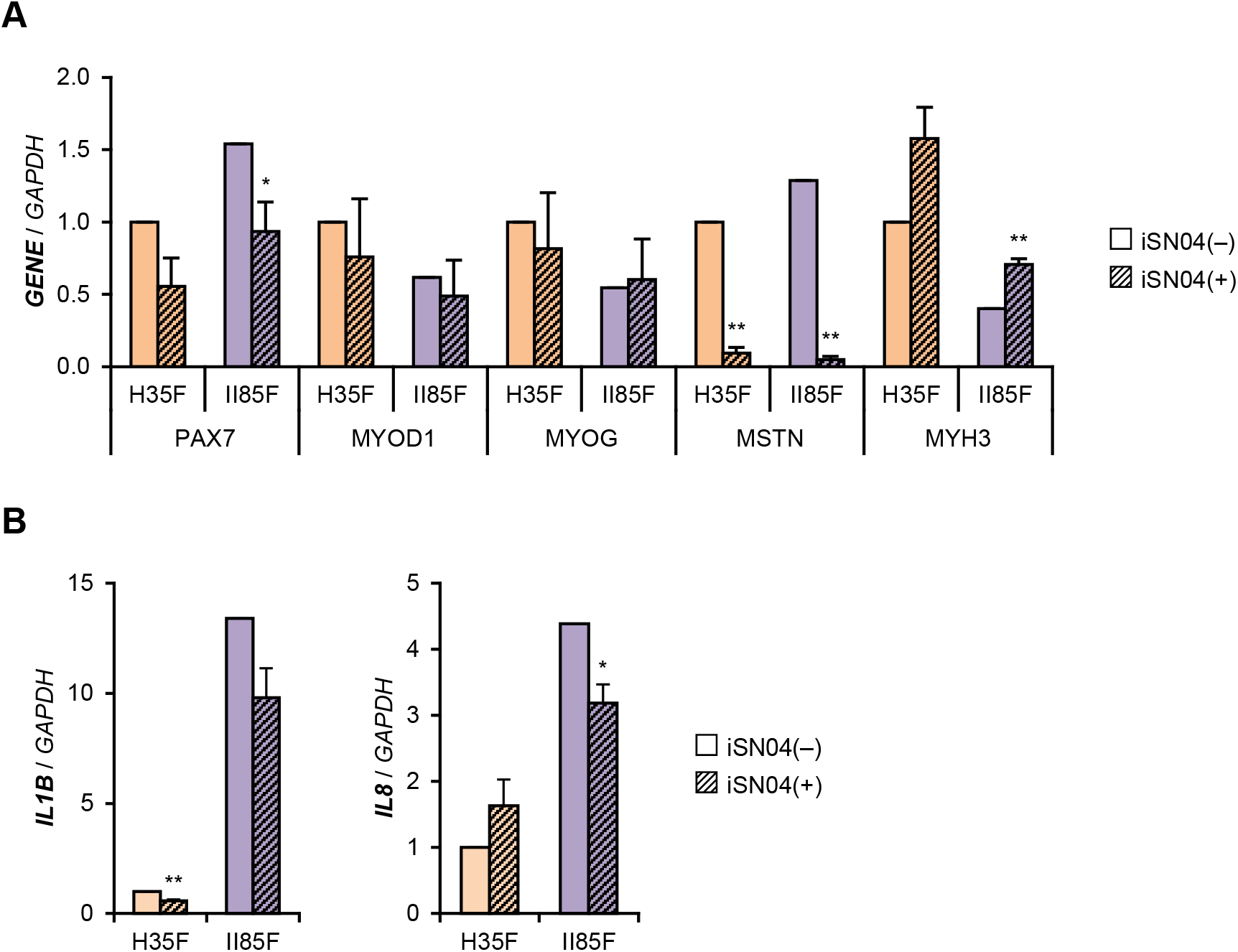
iSN04 suppresses myostatin expression. **(A and B)** qPCR results of gene expression in the H35F and II85F myoblasts differentiated in DIM-NG with iSN04 for two days. Mean value of H35F-iSN04(-) group was set to 1.0 for each gene. * *p* < 0.05, ** *p* < 0.01 vs. iSN04(-) in each myoblast (Student’s *t* test). *n* = 3.

### myoDN Recovers the Myoblast Differentiation Impaired by Excessive Glucose

The DM myoblasts used in this study were isolated from elderly patients (68, 81, 85, and 89-yo) whose ages were significantly higher than those of the healthy subjects (26, 35, and 35-yo) (*p* < 0.01; Student’s *t*-test). Aging is a factor that compromises myoblast differentiation (Brack et al., 2007). To investigate the impact of DM without aging on myogenesis, we cultured and differentiated myoblasts in a high glucose concentration mimicking hyperglycemia. C2C12 cells maintained in high-glucose media exhibited a decreased ratio of MHC^+^ cells and myotubes (Figure 5A). qPCR revealed that high-glucose culture significantly induced *Mstn* and suppressed *Myog* and *Myh3* expression in C2C12 cells on differentiation day 1 (Figure 5B). It is noteworthy that *Il1b* mRNA levels were not elevated by excessive glucose. High-glucose culture also significantly abrogated the myogenesis of H26M and H35F (Figure 5A). These data demonstrated that excessive glucose is an independent factor for the deterioration of myoblast differentiation.

**Figure 5.**
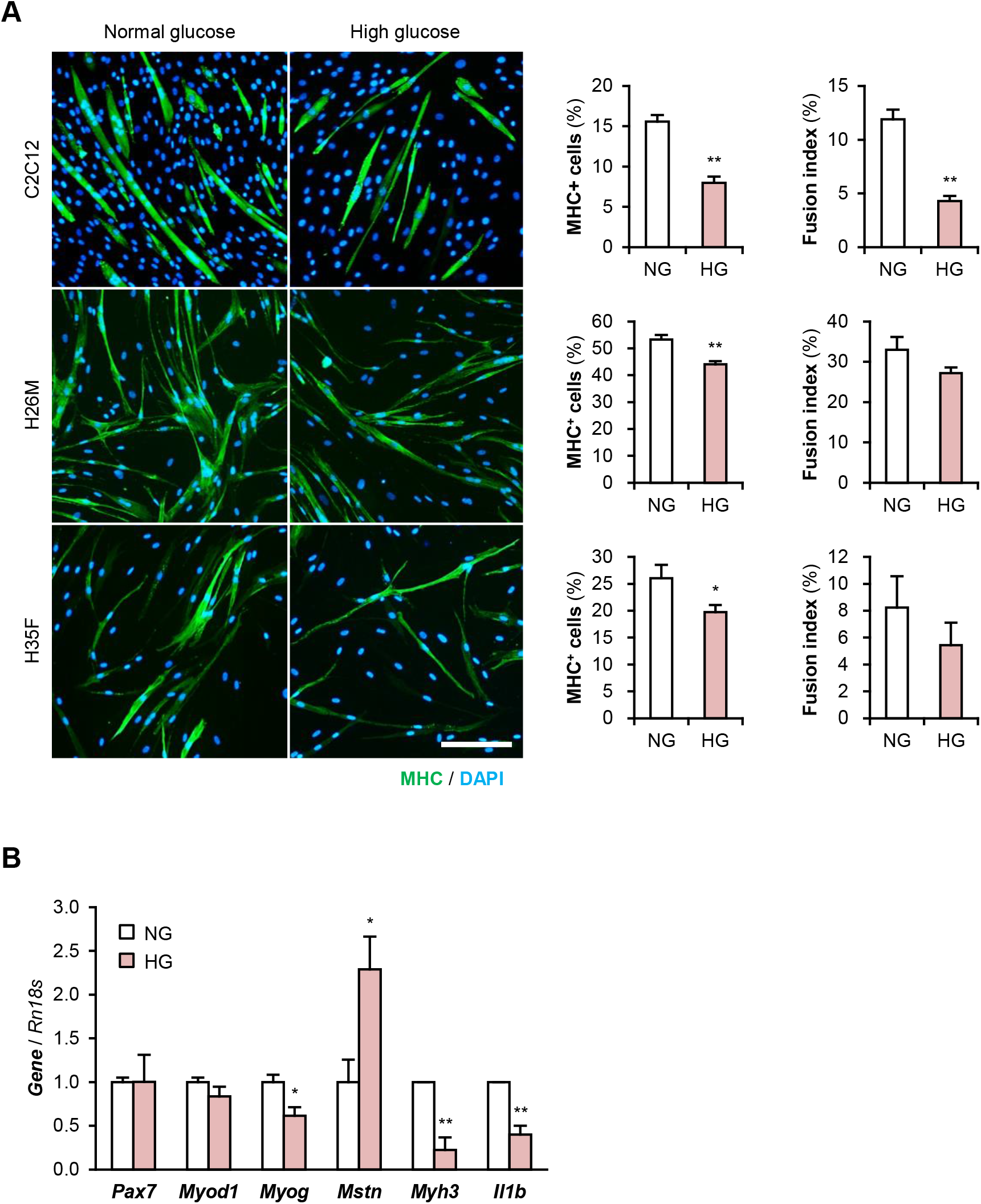
High glucose concentration deteriorates myoblast differentiation. **(A)** Representative immunofluorescent images of the C2C12, H26M, and H35F myoblasts differentiated in DIM-NG or -HG. Scale bar, 200 μm. Ratio of MHC^+^ cells and multinuclear myotubes were quantified. * *p* < 0.05, ** *p* < 0.01 vs. NG (Student’s *t* test). *n* = 4-6. **(B)** qPCR results of myogenic gene expression in the C2C12 cells differentiated in DIM for 1 day. Mean value of NG group was set to 1.0 for each gene. * *p* < 0.05, ** *p* < 0.01 vs. NG (Student’s *t* test). *n* = 3.

Importantly, iSN04 treatment significantly recovered myogenic differentiation and myotube formation in C2C12 cells exposed to high glucose concentrations (Figure 6). This result corresponds well with the phenotype of the iSN04-treated T2DM myoblasts, indicating that myoDNs are potential candidates for nucleic acid drugs that activate myoblasts in hyperglycemic patients.

**Figure 6.**
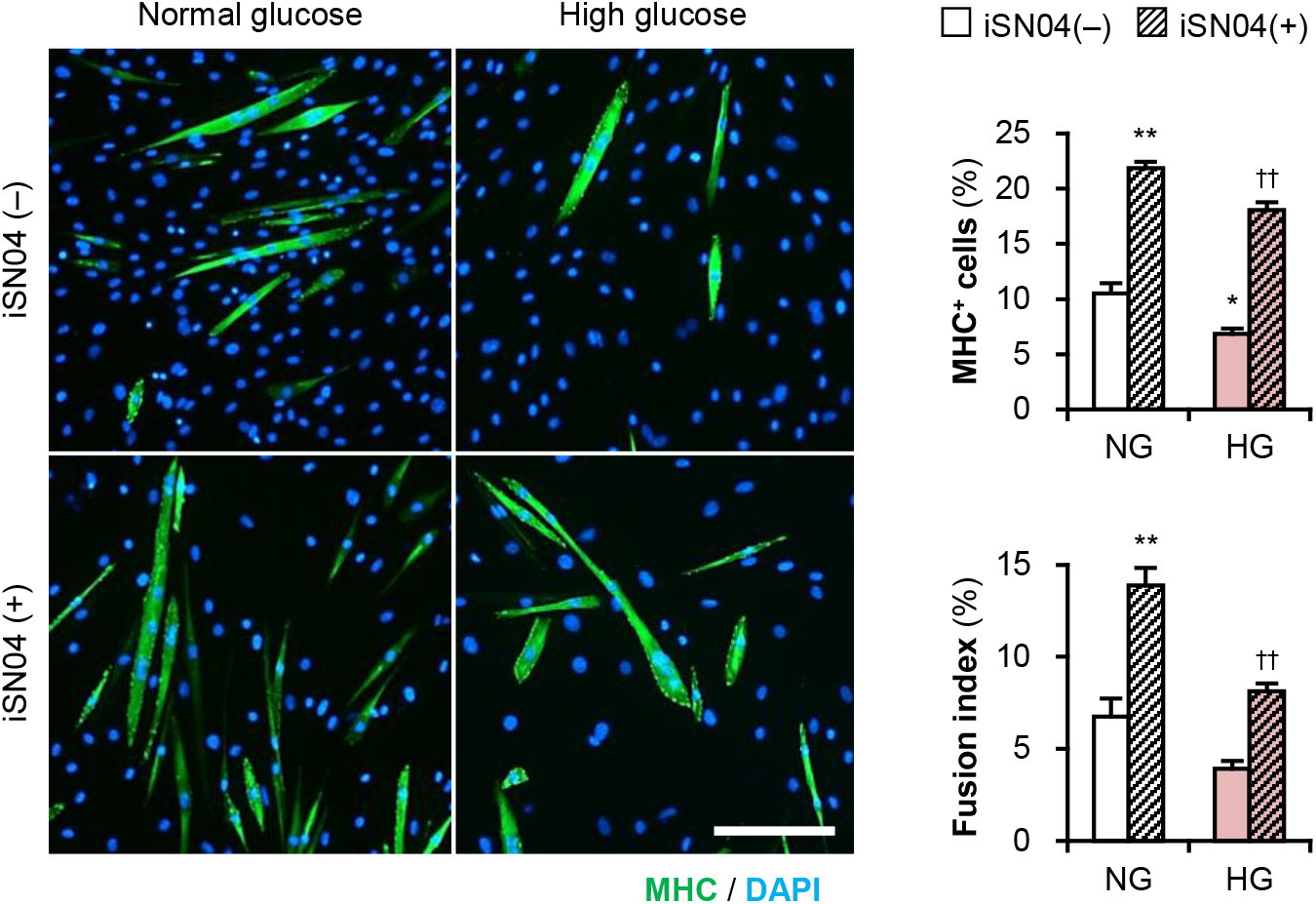
iSN04 recovers the myoblast differentiation impaired by excessive glucose. Representative immunofluorescent images of the C2C12 cells differentiated in DIM with iSN04 for four days. Scale bar, 200 μm. Ratio of MHC^+^ cells and multinuclear myotubes were quantified. * *p* < 0.05, ** *p* < 0.01 vs. NG-iSN04(-); ^††^ *p* < 0.01 vs. HG-iSN04(-) (Tukey-Kramer test). *n* = 4.

### myoDN Recovers the Myoblast Differentiation Impaired by Palmitic Acid

Patients with T2DM are frequently present with hyperlipidemia. Palmatic acid is the most abundant intravital fatty acid, which is involved in insulin resistance and C2C12 cell differentiation (Yang et al., 2013; Saini et al., 2017). To examine the impact of excessive fatty acids on hMBs, H26M and H35M were induced to differentiate in DIM-NG with palmitic acid. In both hMBs, palmitic acid significantly impaired myogenic differentiation and myotube formation (Figure 7A). qPCR showed that palmitic acid decreased the *MYOG/MYOD1* ratio, resulting in lower *MYH3* expression in H35M (Figure 7B). Palmitic acid also upregulated *IL1B* and *IL8* mRNA levels without altering *NFKB1, RELA*, and *TNF*(Figure 7C), which recapitulated the phenotype of T2DM myoblasts. These results indicate that excessive fatty acids can inhibit myoblast differentiation by inducing inflammatory cytokines.

**Figure 7.**
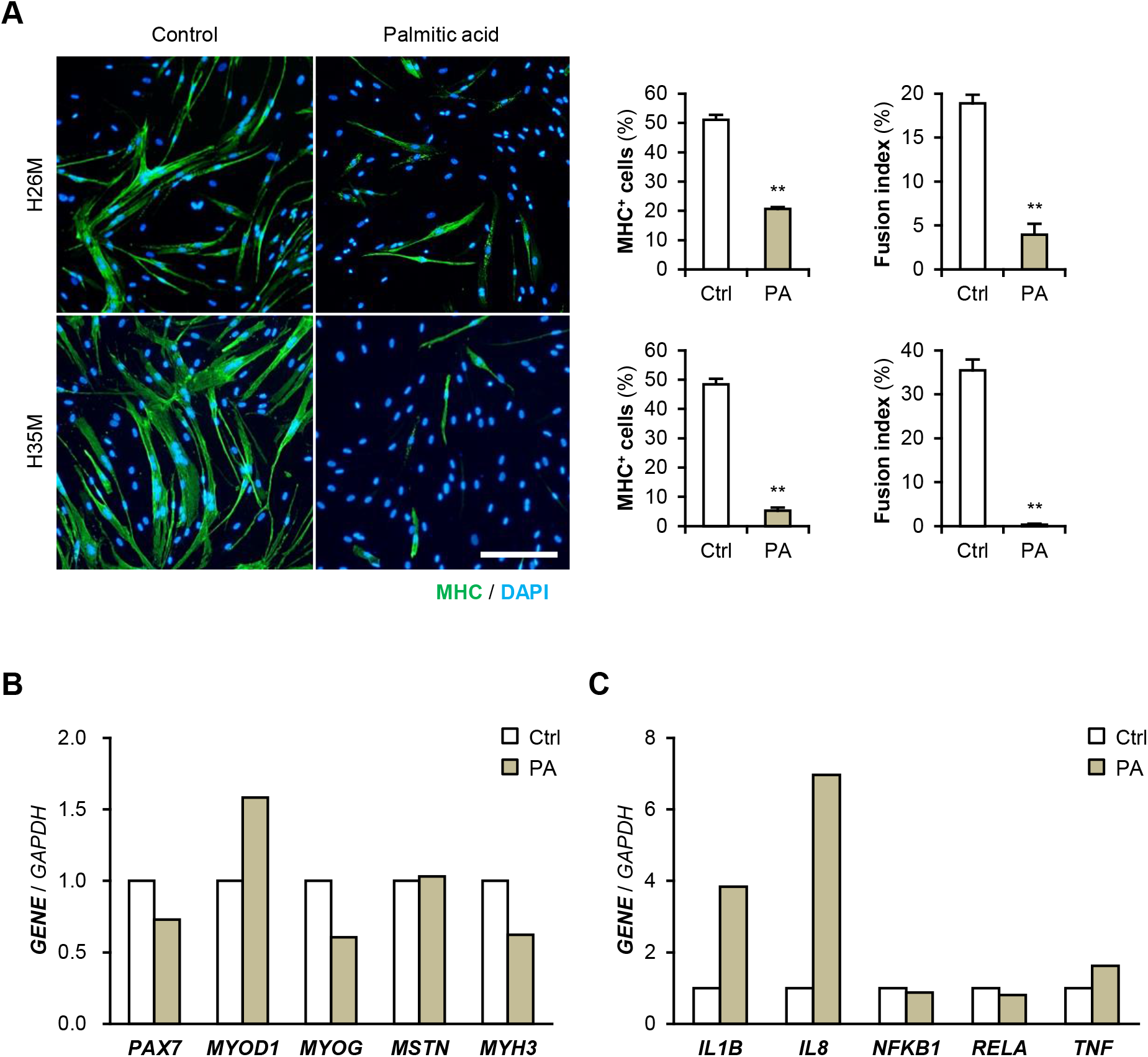
Palmitic acid deteriorates myoblast differentiation. **(A)** Representative immunofluorescent images of the H26M and H35F myoblasts differentiated in DIM-NG with palmitic acid (PA) for two days. Scale bar, 200 μm. Ratio of MHC^+^ cells and multinuclear myotubes were quantified. ** *p* < 0.01 vs. control (Student’s *t* test). *n* = 4-6. **(B and C)** qPCR results of gene expression in the H35M myoblasts differentiated in DIM-NG with palmitic acid for two days. Mean value of control group was set to 1.0 for each gene. *n* = 1.

iSN04 treatment significantly improved the differentiation into MHC^+^ cells from palmitic acid-treated H35M (Figure 8A). As shown in Figure 8B, iSN04 induced *MYOD1* and *MYOG* expression under basal conditions, but not in the presence of palmitic acid. In contrast, iSN04 significantly reduced *MSTN* mRNA levels regardless of the presence of palmitic acid. iSN04 further suppressed palmitic acid-induced *IL8* transcription. These results show that myoDNs conceivably recover myoblast differentiation attenuated by excessive fatty acids in hyperlipidemic patients.

**Figure 8.**
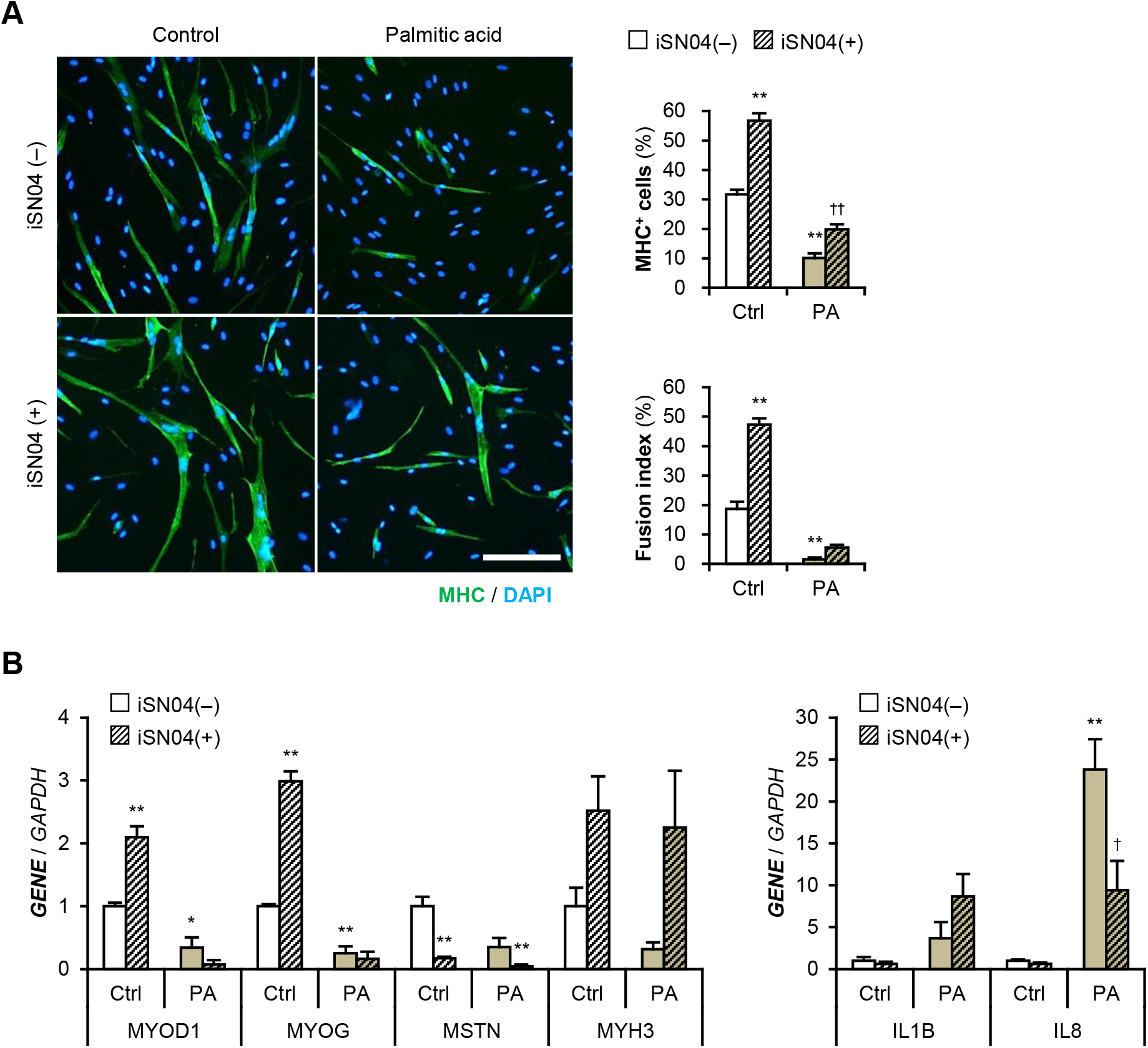
iSN04 recovers the myoblast differentiation impaired by excessive palmitic acid. **(A)** Representative immunofluorescent images of the H35M myoblasts differentiated in DIM-NG with palmitic acid and iSN04 for two days. Scale bar, 200 μm. Ratio of MHC^+^ cells and multinuclear myotubes were quantified. ** *p* < 0.01 vs. control-iSN04(-); ^††^ *p* < 0.01 vs. PA-iSN04(-) (Tukey-Kramer test). *n* = 6. **(B)** qPCR results of gene expression in the H35M myoblasts differentiated as in panel (A). Mean value of control-iSN04(-) group was set to 1.0 for each gene. * *p* < 0.05, ** *p* < 0.01 vs. control-iSN04(-); ^†^ *p* < 0.05 vs. palmitic acid-iSN04(-) (Tukey-Kramer test). *n* = 3.

## Discussion

This study provides evidence that the myoDN, iSN04, ameliorates the differentiation of DM myoblasts, and presents a novel therapeutic strategy for sarcopenic obesity. Dysfunction of DM myoblasts is caused by various pathophysiological factors such as inflammation (D’Souza et al., 2013; Teng and Huang, 2019) and it can be one of the reasons for muscle atrophy. A decreased number of satellite cells has been reported in patients with T1DM (D’Souza et al., 2016). Our results further showed the impaired myogenic ability of T1DM myoblasts with a delayed shift to myogenin-dominant transcription. A similar attenuation of myogenesis has been reported in T2DM myoblasts (Henriksen et al., 2017; Henriksen et al., 2019). The T2DM myoblasts used in this study exhibited a diminished ratio of MyoD/Pax7 and elevated levels of IL-1β and IL-8, which may contribute to the incompetent differentiation. As many patients with T2DM are accompanied by hyperlipidemia in addition to hyperglycemia, surplus glucose and fatty acids are considered the major molecules that interfere with myoblast differentiation. In this study, excessive glucose upregulated myostatin and downregulated myogenin and MHC in C2C12 cells, which is consistent with that reported in previous studies (Grzelkowska-Kowalczyk et al., 2013; Jeong et al., 2013). Similarly, high-glucose culture inhibited myogenesis of plural healthy hMBs. This demonstrates that glucose is an independent factor for myoblast dysfunction, which modulates myogenic gene expression. However, high-glucose culture did not induce IL-1β.

Palmitic acid inhibits myokine expression and C2C12 cell differentiation (Yang et al., 2013; Saini et al., 2017). We showed that palmitic acid abrogates the differentiation of healthy hMBs by upregulating IL-1β and IL-8. IL-1β is known to inhibit IGF-induced myogenin expression and myogenesis (Broussard et al., 2004). IL-8 is a chemokine that contributes to insulin resistance in patients with T2DM (Kim et al., 2006; Samaras et al., 2010) and is also a myokine released from skeletal muscle cells. Insulin-resistant human myotubes secrete higher levels of IL-8 (Bouzakri et al., 2011). The role of IL-8 in myoblast differentiation remains controversial. IL-8-neutralizing antibody impairs the differentiation of hMBs (Polesskaya et al., 2016). In contrast, IL-8 treatment decreases the myogenin/MyoD ratio and embryonic MHC expression in rat myoblasts (Milewska et al., 2019). An appropriate level of IL-8 is important for normal myogenesis. Perturbation of IL-8 in T2DM and palmitic acid-cultured myoblasts may be linked to deteriorated differentiation. The mechanism of IL induction in myoblasts remains unclear. NF-κB p65 and TNF-α have been reported to be elevated in T2DM myoblasts (Green et al., 2011). However, in this study, mRNA levels of these genes were not altered by T2DM or palmitic acid. The signaling pathway of fatty acid-dependent IL induction needs to be clarified in further studies.

This study proved that iSN04 can recover the deteriorated myogenesis of DM myoblasts, in addition to facilitating the differentiation of healthy myoblasts. Although myoDNs, including iSN04, can be potential drug seeds for sarcopenic obesity, the effect of iSN04 should be established using extensive patient-derived myoblasts for clinical application. For instance, the sensitivities to iSN04 were individually different among hMBs. iSN04 is incorporated into the cytoplasm and physically interacts with and interfere with a multifunctional phosphoprotein, nucleolin (Shinji et al., 2021). Nucleolin (*NCL*) mRNA levels were similar among the hMBs used in this study (Supplementary Figure S7A), and subcellular localization of nucleolin was not different between insensitive H26M and sensitive H35F throughout differentiation (Supplementary Figure S7B). Post-translational phosphorylation or glycosylation is indispensable for nucleolin function (Barel et al., 2001; Losfeld et al., 2009). This suggests that the modification of nucleolin may vary among individuals and may be related to iSN04 sensitivity. The precise role of nucleolin in myoblasts remains unclear. One study reported that a moderate knockdown of nucleolin by miR-34b promotes myoblast differentiation (Tang et al., 2017). We found that iSN04 serves as a nucleolin antagonist and increases p53 protein levels to promote myoblast differentiation (Shinji et al., 2021) because nucleolin binds to p53 mRNA to inhibit its translation (Takagi et al., 2005; Chen et al., 2012). However, inhibition of p53 translation is considered to be a part of the multifunction of nucleolin in myoblasts. In cancer cells, nucleolin competitively interacts with NF-κB essential modulator (NEMO), resulting in the downregulation of NF-κB activity. The established nucleolin aptamer AS1411 forms the NEMO-nucleolin-AS1411 complex to block the transcriptional activity of NF-κB (Girvan et al., 2006). We have already confirmed that AS1411 promotes myoblast differentiation as well as iSN04 (Shinji et al., 2021). Thus, iSN04 possibly inhibits NF-κB activity by associating the NEMO-nucleolin-iSN04 complex. NF-κB has been known to impair myogenesis by upregulating Pax7 and myostatin (Wang et al., 2007; He et al., 2013; Ono and Sakamoto, 2017). Inactivation of NF-κB by iSN04 can be assumed to downregulate myostatin and IL-8 in T2DM and palmitic acid-cultured myoblasts. Investigation of anti-inflammatory effects of iSN04 and AS1411 in myoblasts should be an important subject to reveal their action mechanism and to establish the myoDNs as nucleic acid drugs for sarcopenic obesity.

## Conclusion

The differentiation abilities of myoblasts deteriorated with dysregulation of myogenic and inflammatory gene expression due to DM, glucose, or palmitic acid. A myoDN, iSN04, recovered impaired myogenesis by modulating gene expression, especially by decreasing myostatin and IL-8. iSN04 could be a potential drug candidate for sarcopenic obesity by directly targeting myoblasts.

## Author Contributions

TT and SN designed the study. TT wrote the manuscript. SN performed experiments and analyses. SY and TS provided the materials and supervised the study.

