## Supplementary Materials for "Myogenetic oligodeoxynucleotide (myoDN) recovers the differentiation of skeletal muscle myoblasts deteriorated by diabetes mellitus"

### Supplementary Material

**Supplementary Table S1.** qPCR primer sequences for human genes.

| Gene | Sequence (5'-3') | Reference |
|---|---|---|
| *ACVR2B* | GGCTGCTGGCTAGATGACTT<br>AAGCGTTCGTTGCAGAAGTT | Senanayake et al., 2012 |
| *CPT2* | AGCCTCTCTTGAATGATGGCC<br>GATAGGTACATATCAAACCAGGG | Boufroura et al., 2018 |
| *FASN* | CTTCCGAGATTCCATCCTACGC<br>TGGCAGTCAGGCTCACAAACG | Sun et al., 2018 |
| *FBXO32* | CCCAAGGAAAGAGCAGTATGGAGA<br>GGGTGAAAGTGAAACGGAGCA | D'Hulst et al., 2013 |
| *FST* | TGCTCTGCCAGTTCATGG<br>CTTGACGGAGCCAGCAGT | Cheng et al., 2014 |
| *GAPDH* | TGTCAAGCTCATTTCCTGGTA<br>GTGAGGGTCTCTCTCTTCCTCTTGT | Shinji et al., 2020 |
| *IFNG* | AGGGAAGCGAAAAAGGAGTCA<br>GGACAACCATTACTGGGATGCT | Chege et al., 2010 |
| *IL1B* | TCCCCAGCCCTTTTGTTGA<br>TTAGAACCAAATGTGGCCGTG | Sjolinder et al., 2012 |
| *IL6* | CGGGAACGAAAGAGAAGCTCTA<br>GAGCAGCCCCAGGGAGAA | Grosse et al., 2012 |
| *IL8* (*CXCL8*) | TGGCAGCCTTCCTGATTTCT<br>GGGTGGAAAGGTTTGGAGTATG | Grosse et al., 2012 |
| *IRS1* | TATGCCAGCATCAGTTTCCA<br>TTGCTGAGGTCATTTAGGTCTT | Zhao et al., 2017 |
| *IRS2* | TTCTTGTCCCACCACTTGAA<br>CTGACATGTGACATCCTGGTG | Zhao et al., 2017 |
| *MSTN* | CTACAACGGAAACAATCATTACCA<br>GTTTCAGAGATCGGATTCCAGTAT | Bathgate et al., 2018 |
| *MYF5* | AGAACTACTATAGCCTGCCGG<br>ATCTGTGGCATATACATTTGATACATCA | Zibat et al., 2010 |

| | | |
|---|---|---|
| *MYH3* | GGACAGGAAGAATGTGCTGAGATT<br>GCCTCTTGTAGGACTTGACTTTCAC | Shinji et al., 2021 |
| *MYOD1* | TGCTCCGACGGCATGATGGAC<br>TCGACACCGCCGCACTCT | Shinji et al., 2021 |
| *MYOG* | AACCCAGGGGATCATCTGCTCAC<br>GTTGGGCATGGTTTCATCTGGGAAG | Shinji et al., 2021 |
| *NCL* | ATTGGTAGCAACTCCTGGTAAG<br>CACTGTCATCATCCTCCTCTTC | Shinji et al., 2021 |
| *NFKB1* | CACAAGGCAGCAAATAGACG<br>GAGTTAGCAGTGAGGCACCA | Zhao et al., 2018 |
| *PAX3* | AGGAAGGAGGCAGAGGAAAG<br>CAGCTGTTCTGCTGTGAAGG | Sato et al., 2019 |
| *PAX7* | GACCCCTGCCTAACCACATC<br>GTCTCCTGGTAGCGGCAAAG | Shinji et al., 2021 |
| *RELA* | CGCTTCTTCACACACTGGATTC<br>ACTGCCGGGATGGCTTCT | Grosse et al., 2012 |
| *SLC2A4* | CATCCTGATGACTGTGGCTC<br>TCTCATCTGGCCCTAAATACT | Armoni et al., 2005 |
| *SREBF1* | TCCCAGCCCCTCAGATACCAC<br>CCCATTGAGCAGCCAGACCAC | Yang et al., 2019 |
| *SREBF2* | CCCTCACCACCCCTATCCAGA<br>CTCTTGCCCCATCATTACAGG | Yang et al., 2019 |
| *TNF* | CCTGCCCCAATCCCTTTATT<br>CCCTAAGCCCCCAATTCTCT | Sjolinder et al., 2012 |
| *TP53* | AGGCCTTGGAACTCAAGGAT<br>CCCTTTTTGGACTTCAGGTG | Henriksen et al., 2017 |
| *TRIM63* | AAACAGGAGTGCTCCAGTCGG<br>CGCCACCAGCATGGAGATACA | D'Hulst et al., 2013 |
| *TXNIP* | GGCTAAAGTGCTTTGGATGC<br>AGGTCTCATGATCACCATCTCA | Houshmand-Oeregaard et al., 2017 |

**Supplementary Table S2.** qPCR primer sequences for murine genes.

| | | |
|---|---|---|
| *Il1b* | TTGACGGACCCCAAAAGATG<br>CAGGACAGCCCAGGTCAAA | Qu et al., 2009 |
| *Mstn* | ATGGCCATGATCTTGCTGTA<br>CCTTGACTTCTAAAAAGGGATTCA | Han et al., 2010 |
| *Myh3* | CACCTGGAGAGGATGAAGAAGAA<br>AGGACTTGACTTTCACTTGGAGTTTATC | This study |
| *Myod1* | GATGGCATGATGGATTACAGCGGC<br>GTGGAGATGCGCTCCACTATGCTG | This study |
| *Myog* | CCCTATTTCTACCAGGAGCCCCAC<br>GCGCAGGATCTCCACTTTAGGCAG | Watanabe et al., 2011 |
| *Pax7* | CGCGTCCAGGTCTGGTTCAGTAAC<br>GTACTGTGCTGCCTCCATCTTGGG | This study |
| *Rn18s* | CGCACGGCCGGTACAGTGAAACTG<br>CACCCGTGGTCACCATGGTAGGCA | Nihashi et al., 2019 |

#### Supplementary Figure Legends

**Supplementary Figure S1.** The hMBs used in this study. Representative images of the hMBs maintained in hMB-GM-NG. Scale bars, 500 µm (×40) and 50 µm (×200).

**Supplementary Figure S2.** Attenuated myogenic differentiation of DM myoblasts. Ratio of $MHC^{+}$ cells and multinuclear myotubes of the hMBs differentiated in DIM-NG for 0, 2, and 4 days. Bars indicate mean values of each group.

**Supplementary Figure S3.** qPCR results of muscle atrophic gene expression in the hMBs differentiated in DIM-NG for 0, 2, and 4 days. Bars indicate mean values of each group. The mean value of healthy myoblasts at day 0 was set to 1.0 for each gene.

**Supplementary Figure S4.** qPCR results of metabolic gene expression in the hMBs differentiated in DIM-NG for 0, 2, and 4 days. Bars indicate mean values of each group. The mean value of healthy myoblasts at day 0 was set to 1.0 for each gene.

**Supplementary Figure S5.** qPCR results of inflammatory gene expression in the hMBs differentiated in DIM-NG for 0, 2, and 4 days. Bars indicate mean values of each group. The mean value of healthy myoblasts at day 0 was set to 1.0 for each gene.

**Supplementary Figure S6.** iSN04 promotes the differentiation of H26M myoblasts in GM. Representative immunofluorescent images of the H26M differentiated in hMB-GM-NG with 10 µM iSN04 for 2 days. Scale bar, 200 µm. Ratio of $MHC^+$ cells and multinuclear myotubes were quantified. ** $p < 0.01$ vs control (Student's *t* test). $n = 6$.

**Supplementary Figure S7.** Expression and localization of nucleolin in the hMBs used in this study. **(A)** qPCR results of *NCL* expression in the hMBs differentiated in DIM-NG for 0, 2, and 4 days. Bars indicate mean values of each group. The mean value of healthy myoblasts at day 0 was set to 1.0 for each gene. **(B)** Representative immunofluorescent images of the hMBs differentiated in hMB-DIM-NG. Scale bar, 50 µm.

**Figure S1**

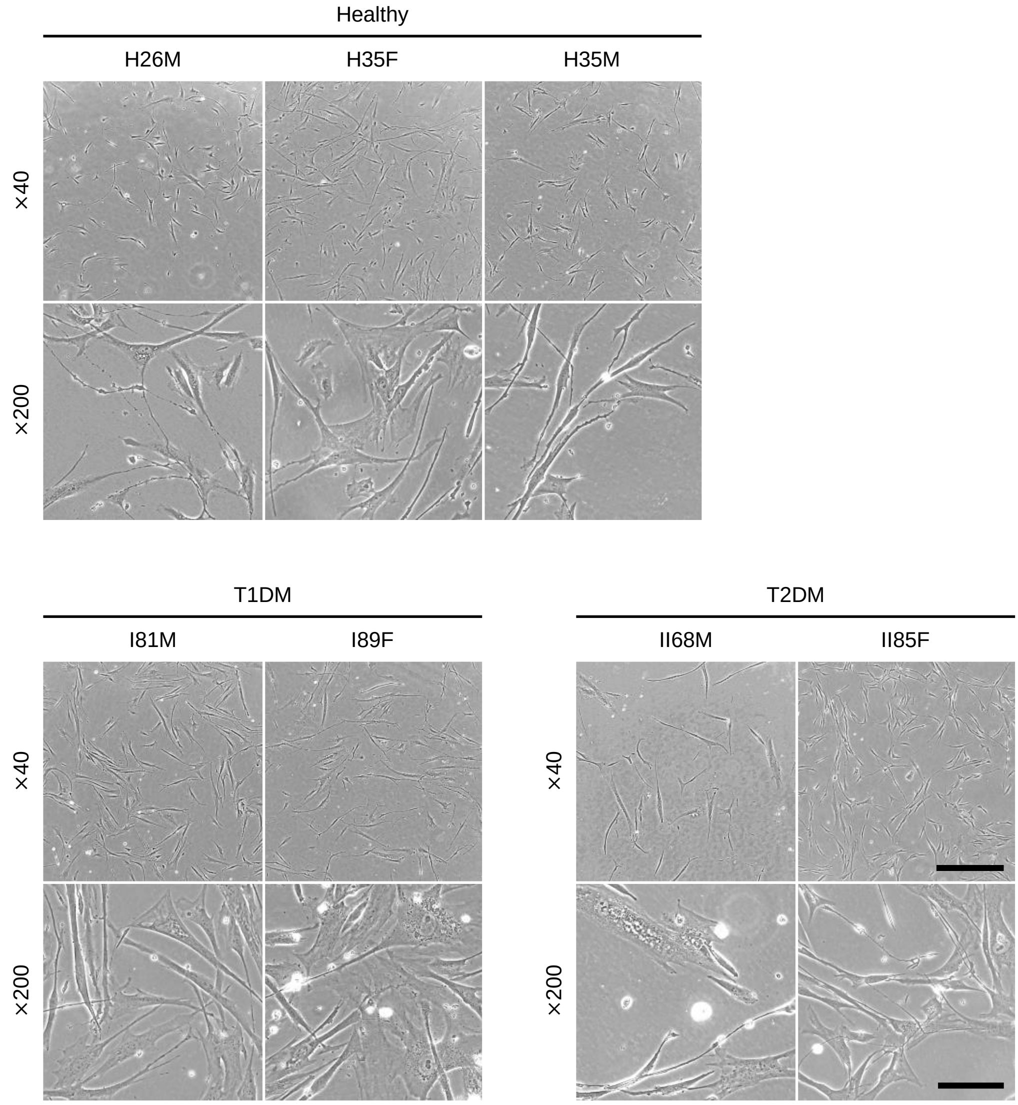
Healthy
H26M
H35F
H35M
×40
×200
T1DM
I81M
I89F
T2DM
II68M
II85F
×40
×200
×40
×200

#### Figure S2

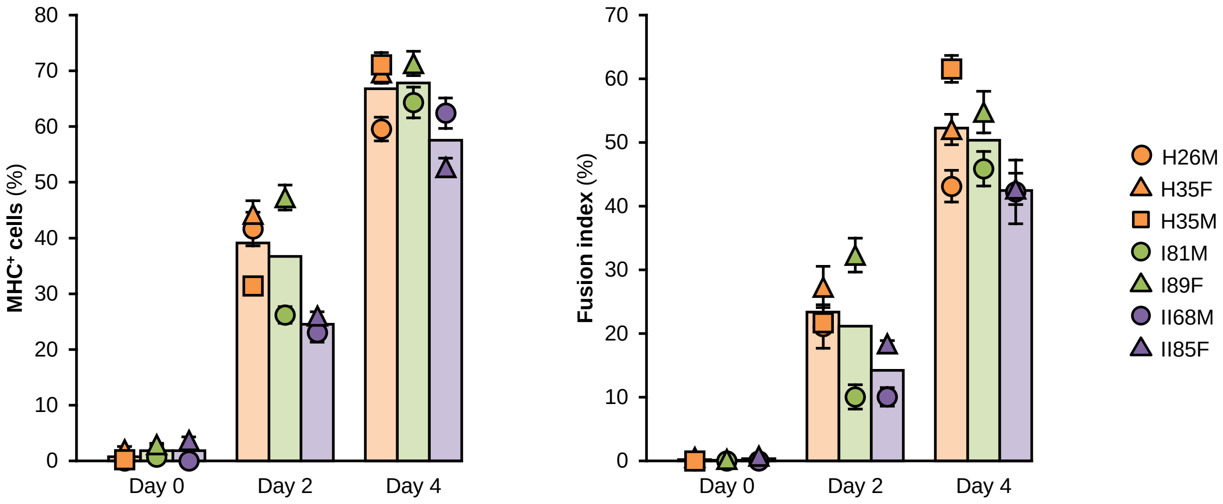

MHC⁺ cells (%)
80
70
60
50
40
30
20
10
0
Day 0
Day 2
Day 4
Fusion index (%)
70
60
50
40
30
20
10
0
Day 0
Day 2
Day 4
H26M
H35F
H35M
I81M
I89F
II68M
II85F

**Figure S3**

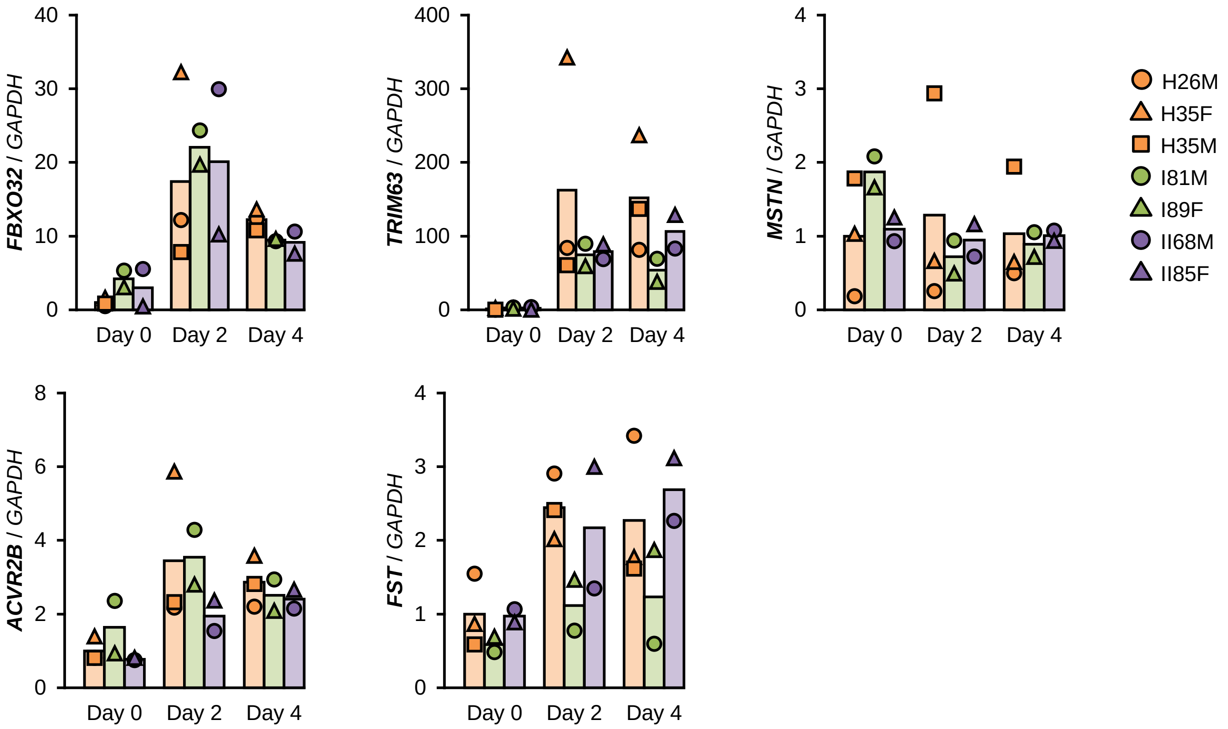

FBXO32 / GAPDH
TRIM63 / GAPDH
MSTN / GAPDH
ACVR2B / GAPDH
FST / GAPDH
Day 0
Day 2
Day 4
H26M
H35F
H35M
I81M
I89F
II68M
II85F

**Figure S4**

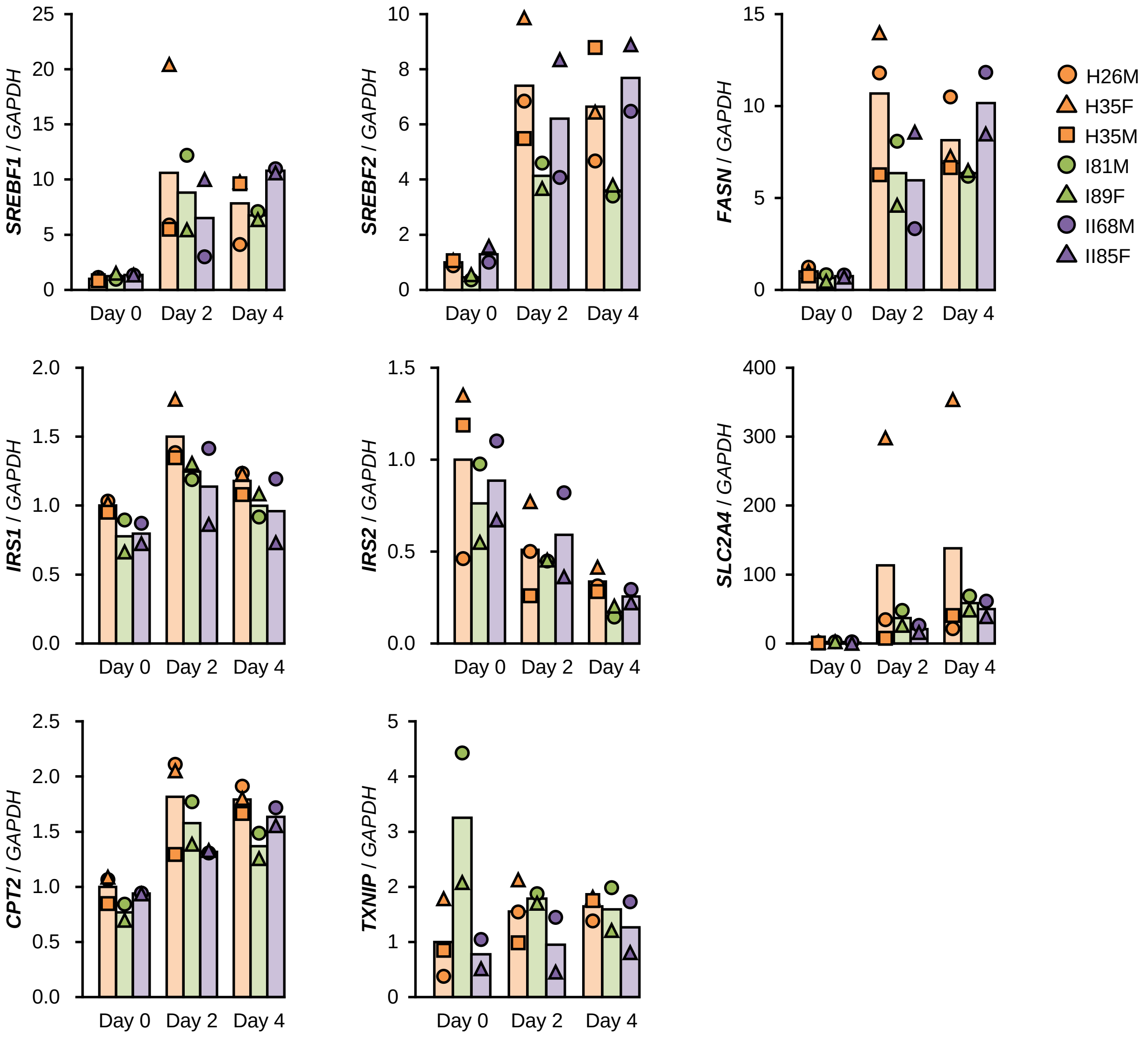

SREBF1 / GAPDH
SREBF2 / GAPDH
FASN / GAPDH
IRS1 / GAPDH
IRS2 / GAPDH
SLC2A4 / GAPDH
CPT2 / GAPDH
TXNIP / GAPDH
Day 0
Day 2
Day 4
H26M
H35F
H35M
I81M
I89F
II68M
II85F

### Figure S5

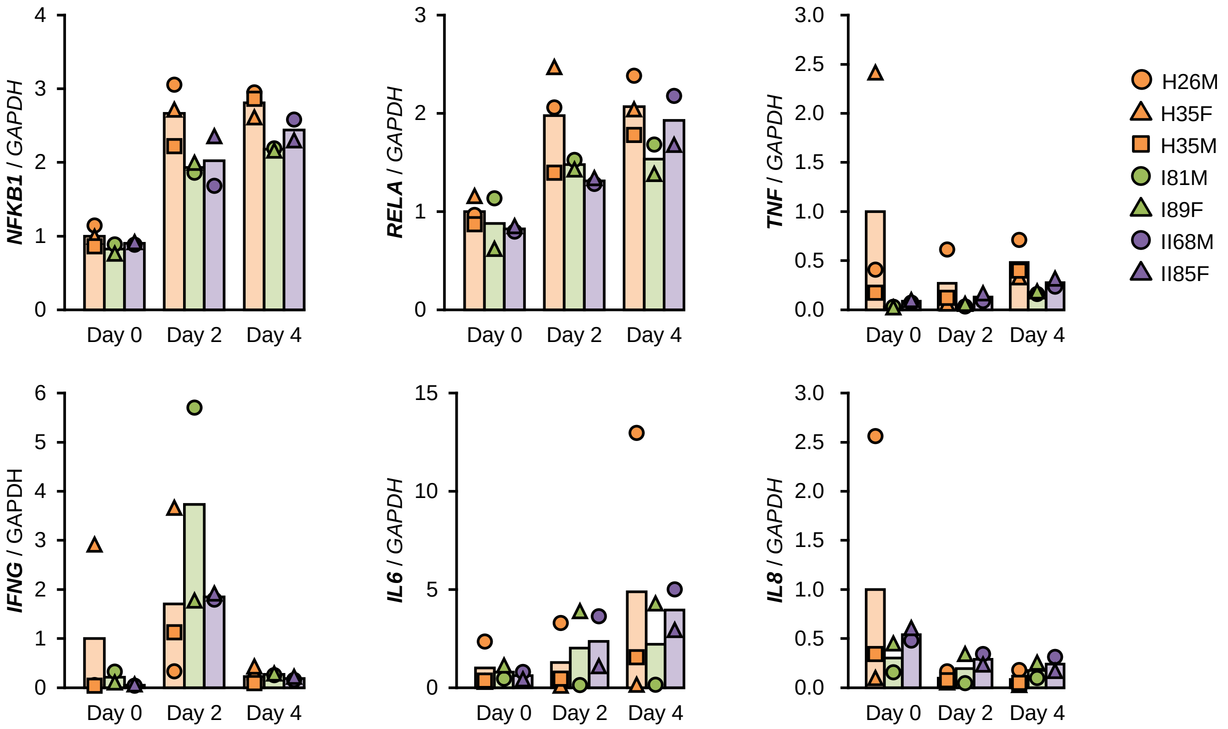
NFKB1 / GAPDH
RELA / GAPDH
TNF / GAPDH
IFNG / GAPDH
IL6 / GAPDH
IL8 / GAPDH
Day 0
Day 2
Day 4
H26M
H35F
H35M
I81M
I89F
II68M
II85F

### Figure S6

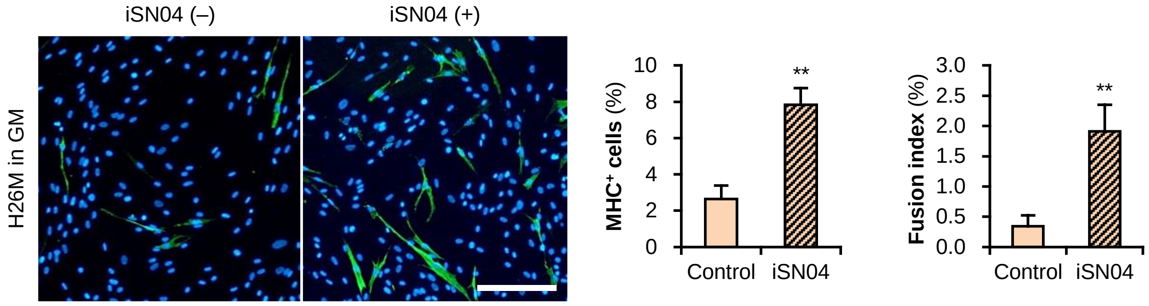

iSN04 (–)
iSN04 (+)
H26M in GM
MHC⁺ cells (%)
10
8
6
4
2
0
**
Control
iSN04
Fusion index (%)
3.0
2.5
2.0
1.5
1.0
0.5
0.0
**
Control
iSN04

**Figure S7**

A

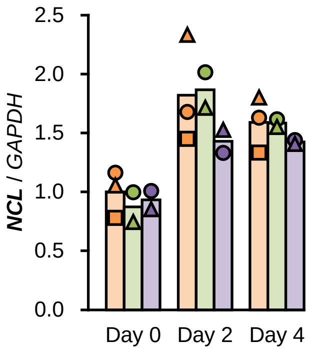

2.5
2.0
1.5
1.0
0.5
0.0
NCL / GAPDH
Day 0
Day 2
Day 4


B

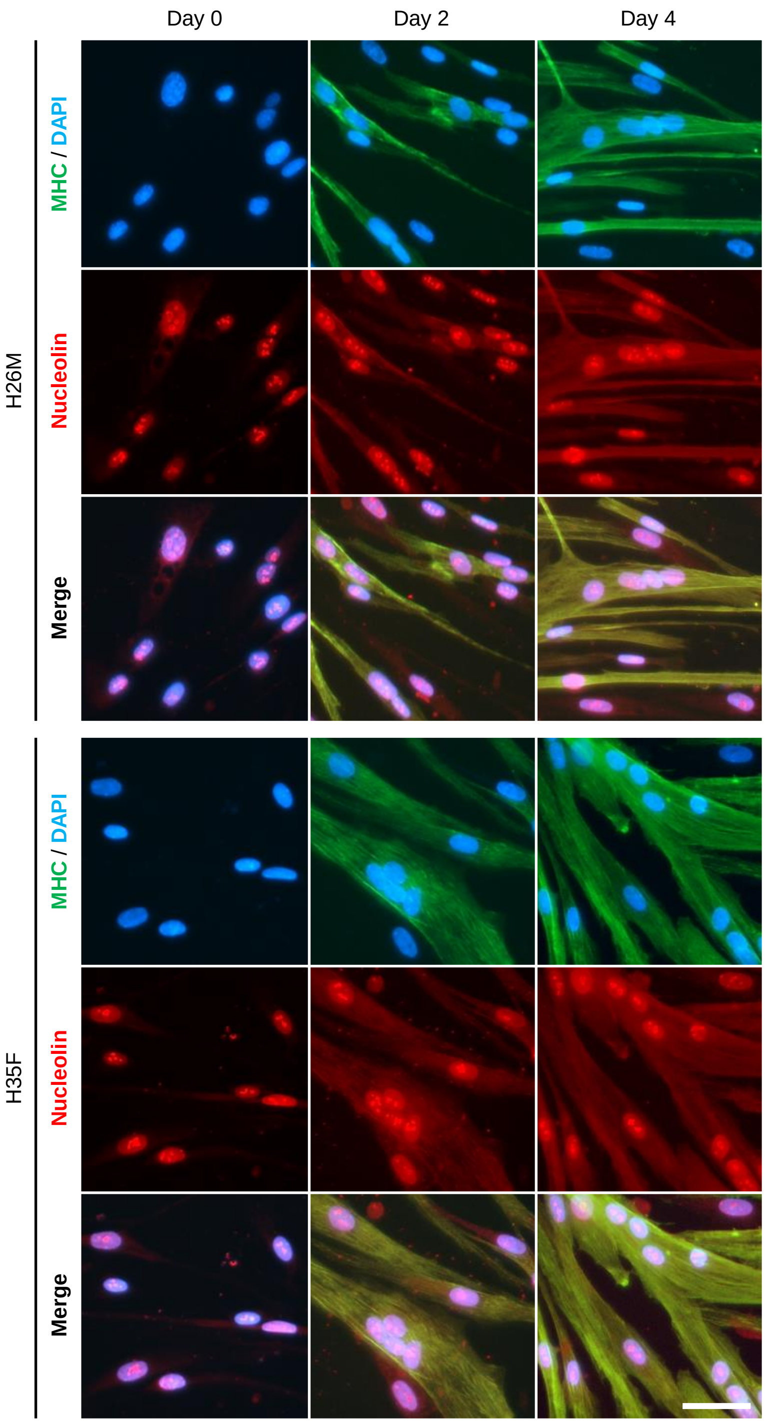

Day 0
Day 2
Day 4
H26M
MHC / DAPI
Nucleolin
Merge
H35F
MHC / DAPI
Nucleolin
Merge
